# Molecular dynamic simulation reveals E484K mutation enhances spike RBD-ACE2 affinity and the combination of E484K, K417N and N501Y mutations (501Y.V2 variant) induces conformational change greater than N501Y mutant alone, potentially resulting in an escape mutant

**DOI:** 10.1101/2021.01.13.426558

**Authors:** Gard Nelson, Oleksandr Buzko, Patricia Spilman, Kayvan Niazi, Shahrooz Rabizadeh, Patrick Soon-Shiong

**Affiliations:** ImmunityBio, Inc.

## Abstract

Rapidly spreading SARS-CoV-2 variants present not only an increased threat to human health due to the confirmed greater transmissibility of several of these new strains but, due to conformational changes induced by the mutations, may render first-wave SARS-CoV-2 convalescent sera, vaccine-induced antibodies, or recombinant neutralizing antibodies (nAbs) ineffective. To be able to assess the risk of viral escape from neutralization by first-wave antibodies, we leveraged our capability for Molecular Dynamic (MD) simulation of the spike receptor binding domain (S RBD) and its binding to human angiotensin-converting enzyme 2 (hACE2) to predict alterations in molecular interactions resulting from the presence of the E484K, K417N, and N501Y variants found in the South African 501Y.V2 strain – alone and in combination. We report here the combination of E484K, K417N and N501Y results in the highest degree of conformational alterations of S RBD when bound to hACE2, compared to either E484K or N501Y alone. Both E484K and N501Y increase affinity of S RBD for hACE2 and E484K in particular switches the charge on the flexible loop region of RBD which leads to the formation of novel favorable contacts. Enhanced affinity of S RBD for hACE2 very likely underpins the greater transmissibility conferred by the presence of either E484K or N501Y; while the induction of conformational changes may provide an explanation for evidence that the 501Y.V2 variant, distinguished from the B.1.1.7 UK variant by the presence of E484K, is able to escape neutralization by existing first-wave anti-SARS-CoV-2 antibodies and re-infect COVID-19 convalescent individuals.

## Introduction

As many SARS-CoV-2 variants emerge and displace first-wave viruses^1,2^, it is important not only to assess their relative transmissibility, but also their ability to escape antibody neutralization by convalescent antibodies in recovered COVID-19 patients^3^, recombinant neutralizing antibodies (nAbs) developed as therapeutics, or antibodies elicited by first-generation vaccines.

Of great interest are variants that include mutations with the potential to affect the interaction of the SARS-CoV-2 spike receptor binding domain (S RBD) with the host receptor, angiotensin-converting enzyme 2 (ACE2). The binding of the S RBD of SARS-CoV-2, like SARS-CoV before it, to ACE2 initiates infection ^4–7^, thus variants that have a greater binding affinity for ACE2 are likely to be more readily transmissible^8^. Transmissibility goes hand-in-hand with mortality, because even if a variant does not produce a higher rate of morbidity or mortality, the total number of severe cases and death would be expected to increase due to what may be an exponential increase in infections.

The dire consequences of more rapid and widespread infection can further be compounded by a decrease in efficacy of available antibody-based therapeutics and vaccines; and by a loss of protective immunity in persons previously infected with a ‘first wave’ virus. The efficacy of vaccines may be altered if a specific mutation or combination of mutations in a variant results in significant conformational changes that render key regions of S that participate in ACE2 binding ‘unrecognizable’ to antibodies generated in response to a first-generation vaccine. A similar principle is in play for the efficacy of nAbs targeted to the receptor interface ^9,10^ and convalescent sera.

Here, to better understand the risks posed by individual or combined mutations in the ‘second-wave’ variants, we leveraged our *in silico* Molecular Dynamic (MD) simulation capabilities to perform computational analysis of interactions of the S RBD with human ACE2. In our first report, Nelson *et al.*^11^ (in preprint) “*Millisecond-scale molecular dynamics simulation of spike RBD structure reveals evolutionary adaption of SARS-CoV-2 to stably bind ACE2*”, we initially used millisecond-scale MD simulation to simulate free SARS-CoV-2 S RBD based on previously reported structures^12,13^ as well as its molecular interactions with ACE2 and showed S adopts a binding-ready conformation, incurring little entropic penalty during ACE2 interaction. We further revealed areas of high-affinity interaction between S RBD and ACE2 that have a high likelihood of determining binding kinetics.

Here, we utilized the MD simulation methods employed in our first study to investigate what effects mutations found at the S RBD-ACE2 interface in the rapidly spreading South African variant 501Y.V2^14^ - E484K, K417N, and N501Y – have on RBD binding affinity and spike conformation.

## Results

As shown in Figure 1a, the E484K, K417N, and N501Y mutants span the S RBD-ACE2 interface, with the E484K substitution occurring in a highly flexible loop region of the S RBD (Fig. 1c). The N501Y substitution is found in a second region of contact^11^, and the K417N mutation in a region between the two that shows relatively little interaction with ACE2.

**Fig. 1.**
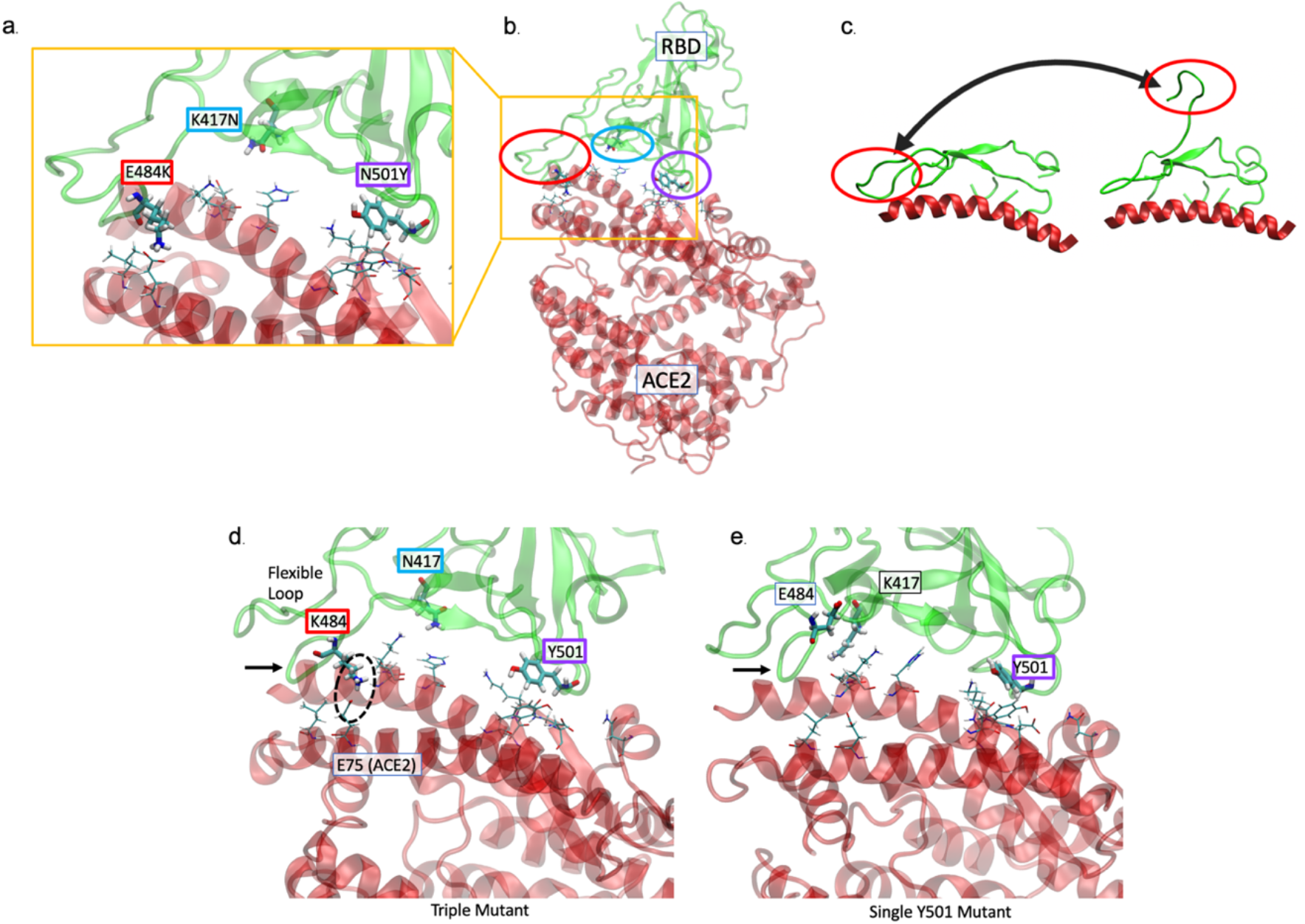
The K484 substitution in the novel South African variant increases affinity of the spike receptor binding domain (S RBD) for ACE2. (a, b) The positions of the E484K (red), K417N (cyan), and N501K (purple) substitutions at the interface of the 501Y.V2 variant S RBD - hACE2 interface are shown. hACE2 residues nearest to the mutated RBD residues are rendered as thin sticks. The E484K mutation is located in a highly flexible loop region of the interface, K417N in a region with lower probability of contact, and N501K at a second point of high-affinity contact. (c) The range of movement available to the loop containing residue 484 is shown by PCA of MD simulation of a first-wave sequence^11,13^. (d) MD simulation performed in the presence of all 3 substitutions reveals the loop region is tightly associated (black arrow) with hACE2. A key contact ion pair is circled. (e) In comparison to K484, when E484 (‘wildtype’) is present with only the Y501 variant, the loop is not as tightly associated (arrow).

### E484K shows enhanced affinity for ACE2

As revealed by our MD simulation, the presence of a lysine residue at position 484 in the presence of the other two substitutions (an asparagine residue at position 417; a tyrosine residue at position 501) resulted in increased affinity at position 484 that can be described as the loop being ‘locked onto’ ACE2 (Fig. 1d) as compared to when a glutamic acid residue is present at position 484 (Fig. 1e) and a tyrosine at 501. This increased affinity of K484 for hACE2 may be due, in part, to the change in net charge on the flexible loop from −1 to +1. This allows the formation of a transient contact ion pair with E75 in hACE2.

### E484K in combination with K417N, and N501Y induces conformational changes in spike

To determine if it is likely the presence of the E484K or N501Y mutations alter the conformation of S RBD, we performed Principal Component Analysis (PCA) for the triple mutant, E484K and N501Y mutants alone, and compared them to the original cryo-EM^13^ structure representing the ‘first wave’ sequence. The PCA plots are shown in Figure 2. While both mutants sample similar conformations, S RBD with K484 alone (Fig. 2b) preferentially adopts conformations similar to the cryo-EM structure described in Wrapp *et al.*^13^. This is similar to the behavior observed for the first-wave strain as described in our initial report ^11^. Y501 alone (Fig. 2c) affects conformational probability, shifting it away from the original structure. For K484, the effect of the contact ion pair is notable as a distinct density in the black dashed circled region of Figure 2b. When all 3 mutations are present (Fig. 2a), other conformations as indicated by the circled regions, are much more likely to occur compared to the original cryo-EM structure, particularly compared to the Y501 alone (Fig. 2c). With the triple mutant, the there is a group of conformations that show good overlap with, and therefore high similarity to, the K484 and WT simulations. There is also a cluster of conformations in a region that is unique to the triple mutant, likely reflecting the influence of the other mutations. Only the WT RBD includes the cryo-EM structure in a region of high probability density (Fig. 2d). The mutated variants adopt novel conformations with varying degrees of distance from the cryo-EM structure. Notably, the single mutants (E484K and N501Y) have a clearly preferred family of conformations while the PCA density of the triple mutant suggests there exists an equilibrium between two distinct states.

**Fig. 2.**
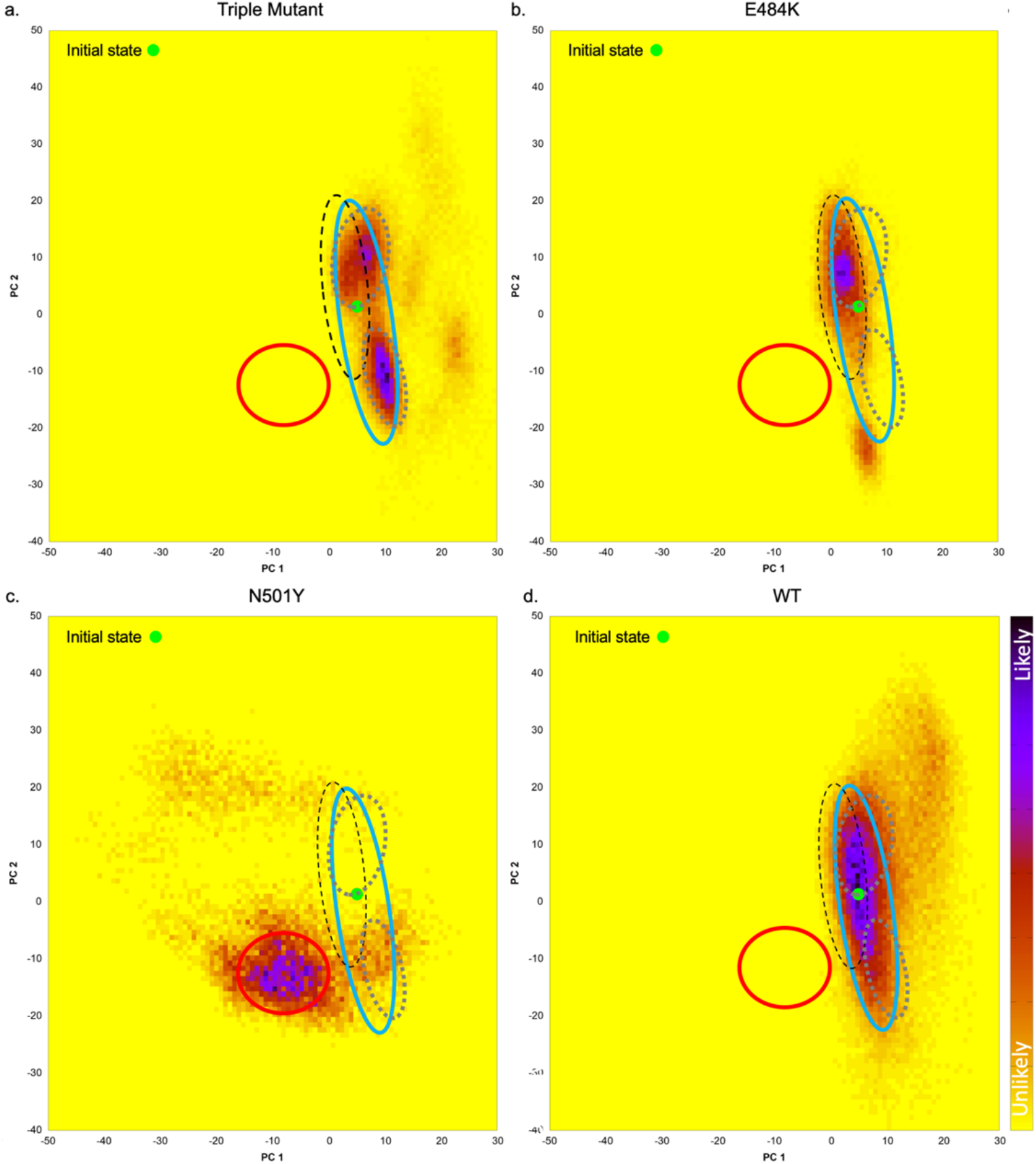
Presence of the E484K mutations alters the conformation of S RBD. Principal component analysis (PCA) plots are shown comparing the accessible conformational space for (a) the triple mutant comprising E484K, K417N, and N501Y; (b) the E484K mutant only; (c) the N501Y mutant only; and (d) the ‘first wave’ sequence to compare the accessible conformational space of each S RBD when bound to hACE2. The green dot indicates the location of the cryo-EM structure, PDB 6M17^13^. The blue circled regions indicate conformations that are similar to the original cryo-EM structure, the red circled region to those found for N501Y, the black dashed circle those found for K484 alone, and the gray dashed circles those found for the triple mutant; with a greater density (purple) indicating a greater probability other conformations occur.

### E484K shows increased contact with hACE2 E75

E484K, whether in the presence of both K417N, and N501Y variants (Fig. 3a), or as the only variant in the presence of K417 and N501 (Fig. 3b) is associated with increased contact between RBD residue 484 and ACE2 E75 compared to either the N501Y mutant alone or the ‘first wave’ (WT) sequence (Fig. 3c and d, respectively). In the presence of either Y501 and WT, E484 shows little contact with hACE2 E75. In addition, The RBD E484K mutant shows increased contact between it and several residue pairs in addition to hACE2 E75.

**Fig. 3.**
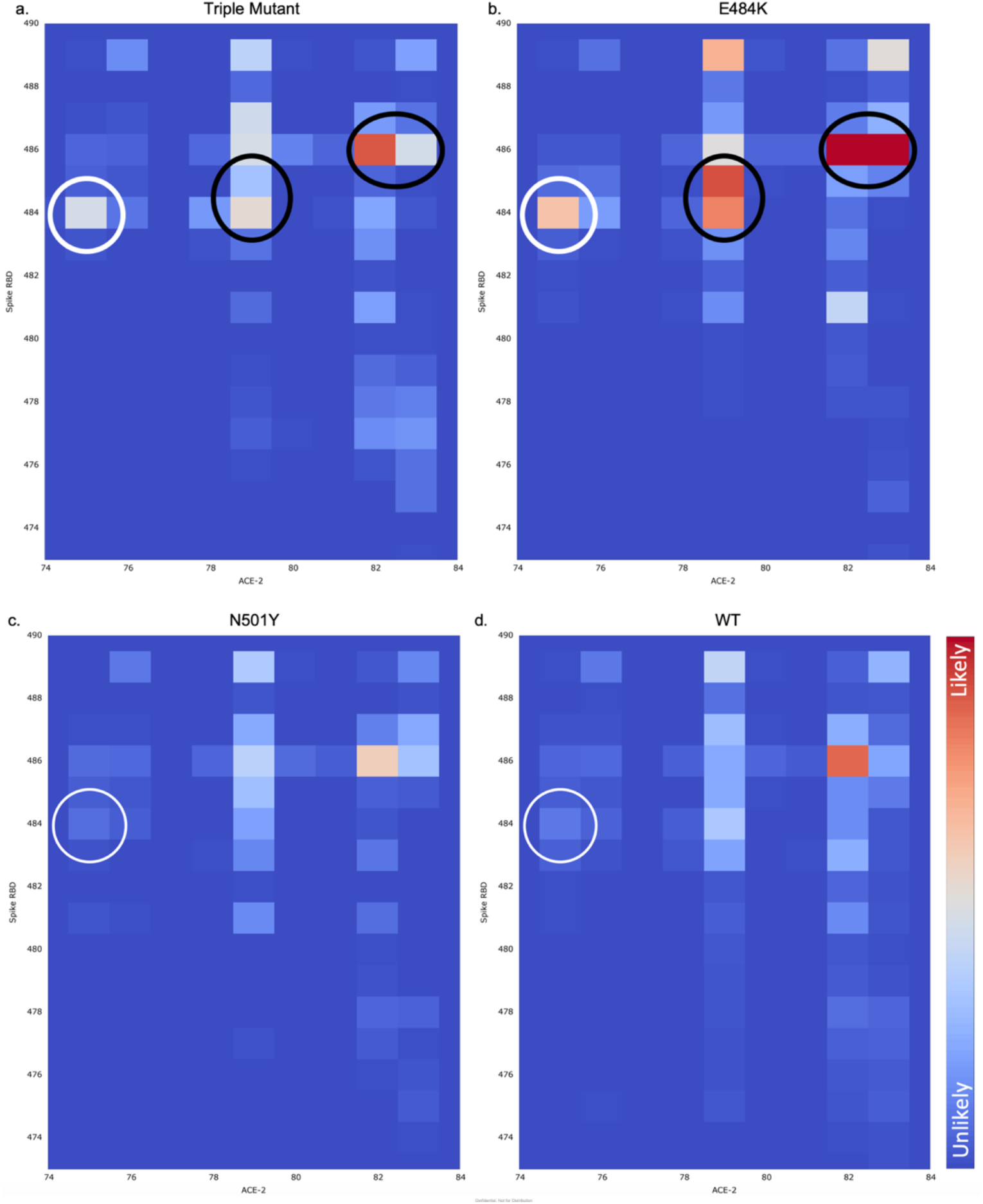
Contact maps for position 484 and hACE2 E75. Contact maps are shown for (a) the triple mutant, (b) E484K, (c) N501Y, and (d) ‘wildtype’ spike. Both the triple mutant and the E484K mutant alone show more contact between RBD position 484 and ACE2 E75 (white circles) than N501Y or wildtype (WT or ‘first wave’ sequence). The RBD E484K mutant also shows increased contact with several other residue pairs (black circles). The color scale is consistent across the plots.

## Discussion

Our MD simulation-based predictions of greater affinity of K484 S RBD for ACE2 as compared to E484 and the greater likelihood of altered conformation as compared to the original structure may represent mechanisms by which the new 501Y.V2 virus variant has been able to replace original viral strains. The enhanced affinity likely accounts for by rapid spread due to greater transmissibility and the likelihood of adoption of conformations not recognized by first-wave antibodies may explain escape of the virus from immunity to the original strain^14^.

The strong contacts seen for only the E484K mutant here suggest that spike may adopt a distinct ‘binding pose’ relative to other conformations. We further posit the differing PCA densities seen for the triple mutant that seem to result in lesser loop interaction with hACE2 as compared to E484K alone may be due to the other mutations. We will investigate this in depth in future studies.

The greater relative risk posed by variants expressing the E484K mutant as compared to the N501Y mutant alone is supported by Andreano *et al.* (in preprint)^15^ who co-incubated what they refer to as ‘authentic’ virus with highly neutralizing COVID-19 neutralizing plasma over several passages and saw the appearance of mutations including the E484K substitution which, in combination with an insertion in the spike N-terminal domain (NTD), led to complete resistance to the convalescent plasma. In contrast, Xie et al.^16^ (in preprint) reported that sera from 20 individuals that received the BNT162b2 mRNA vaccine showed equivalent ability to neutralize both Y501 and N501 SARS-CoV-2.

Additional support for the threat posed by mutations at residue 484 is provided by Greaney *et al.^17^* who undertook an impressive effort to map mutations that affect binding of ten human monoclonal antibodies. By employing a deep mutational scanning method, they found that mutations at residue 484 have a high probability of affecting antibody binding. They further suggested their analytical method provides a tool for design of escape-resistant antibody cocktails to overcome the threat posed by viral evolution. We believe the MD simulation approach used here similarly represents a tool to be used in the arsenal against the continuing pandemic, as it provides insight into the likelihood mutations alone or in combination may have effects that lessen the efficacy of existing therapies or vaccines.

In vaccine design, it has been suggest that the makers of vaccines could keep up with viral evolution by continual alteration of the ‘payload’ - almost all vaccines in development use the spike sequence - to fit currently predominant strains. We suggest vaccines whose efficacies are largely dependent upon humoral responses to the S antigen only are inherently limited by the emergence of novel strains and dependent upon frequent re-design. In contrast, a vaccine that elicits a vigorous T-cell response that is far less subject to changes due to accruing mutations provides a better, more efficient approach to protection. The ideal vaccine would also deliver a second, conserved antigen such as the SARS-CoV-2 nucleocapsid protein, that very likely will elicit humoral and cell-mediated immune responses that will remain effective, even in the face of a rapidly changing virus.

Although not the subject of the investigation described here, we are developing a dual-antigen human adenovirus serotype 5 (Ad5) platform-based vaccine that delivers both a spike protein with a linker to increase cell surface expression and humoral responses (S-Fusion) and the highly antigenic and conserved nucleocapsid (N) protein with a signal sequence (an Enhanced T-cell Stimulation Domain, ETSD) to direct it to subcellular compartments that enhance MHC I and II responses^18^. It is our belief that the vaccine, hAd5 S-Fusion + N-ETSD, due to its ability to elicit cell-mediated in addition to humoral immune responses, as shown in both a rodent model^19^ and non-human primates^20^, offers hope to those regions such as South Africa wherein dangerous variants of SARS-CoV-2 have swept the country.

## Materials and Methods

### System Setup

The WT-ACE2/RBD complex was built from the cryo-EM structure, PDB 6M17 of full-length human ACE2 in the presence of the neutral amino acid transported B^0^AT1 with the S RBD as shown in Yan *et al.*^21^ using RBD residues 336-518 and ACE2 residues 21-614. The appropriate RBD mutations (K417N, E484K and N501Y for the triple mutant and either E484K or N501Y for the single mutants) were introduced and the complex was created using the Amber ff14SB force field ^22^. A thin solvating shell of water was placed around the complex using the RISM program from AmberTools19 ^23^ to determine optimal locations. Bulk waters (40,655, 40,801 and 39,714 waters for the triple mutant, E484K and N501Y, respectively) were added to create a sufficient water box and sodium ions (22, 21 and 23 for the triple mutant, E484K and N501Y, respectively) were added at random locations to neutralize the system.

### Simulation

10 copies of each RBD mutant were minimized, equilibrated and simulated. Minimization occurred in two phases. During the first, the protein and RISM-placed waters were restrained. The second phase minimized the entire system. Dynamics then began and the temperature was ramped from 0 to 300K while restraining the protein and RISM-placed waters. All dynamics used SHAKE restraints on hydrogen-containing bonds and a 2fs timestep. All restraints were then released and the system was equilibrated in the NPT ensemble for 2ns. Finally, the system equilibrated in the NVT ensemble for 100 ns before data collection began. All simulations were performed with the GPU-enabled version of Amber20^23^.

### Principle Component Analysis (PCA)

The backbones of residues at the RBD/hACE2 interface were used for all PCA calculations. Structures were RMSD aligned to the cryo-EM structure. Eigenvectors for the PCA plots were then calculated using the full set of simulations of the triple mutant, E484K and N501Y systems. Simulation structures were projected onto the eigenvectors for each mutation system separately. All calculations were run with cpptraj and plotted using gnuplot.

## Notes

### Competing Interest Statement

All authors are employee or senior management of ImmunityBio Inc. that is developing a vaccine mentioned in the manuscript

